# *In vitro* susceptibility to β-lactam antibiotics and viability of *Neisseria gonorrhoeae* strains producing plasmid-mediated broad- and extended-spectrum β-lactamases

**DOI:** 10.1101/2022.03.11.483876

**Authors:** Ilya Kandinov, Dmitry Gryadunov, Alexandra Vinokurova, Olga Antonova, Alexey Kubanov, Victoria Solomka, Julia Shagabieva, Dmitry Deryabin, Boris Shaskolskiy

**Affiliations:** Center for Precision Genome Editing and Genetic Technologies for Biomedicine, Engelhardt Institute of Molecular Biology, Russian Academy of Sciences, 119991 Moscow, Russia; State Research Center of Dermatovenerology and Cosmetology, Russian Ministry of Health, 107076 Moscow, Russia

**Keywords:** *Neisseria gonorrhoeae*, β-lactamase-producing plasmids, *N. gonorrhoeae* viability, extended-spectrum β-lactamase

## Abstract

*N. gonorrhoeae* strains producing plasmid-mediated broad- and extended-spectrum β-lactamases (ESBLs) have been obtained *in vitro* and studied for their viability and susceptibility to β-lactam antibiotics. The artificial p*bla*_TEM-1_ and p*bla*_TEM-20_ plasmids were constructed by site-directed mutagenesis from a p*bla*_TEM-135_ plasmid of the Toronto/Rio type, which was extracted from the clinical isolate. MIC values were determined for a series of β-lactam antibiotics, including benzylpenicillin, ampicillin, cefuroxime, ceftriaxone, cefixime, cefotaxime, cefepime, meropenem, imipenem, and doripenem. The *N. gonorrhoeae* strain carrying the p*bla*_TEM-20_ plasmid exhibited a high level of resistance to penicillins and II-IV generation cephalosporins (MIC ≥ 2 mg/L) but not to carbapenems (MIC ≤ 0.008 mg/L). However, this strain stopped growing after 6 hours of cultivation. The reduced viability was not associated with the plasmid loss but can be explained by the following factors: the presence of the plasmid itself, which requires additional costs for its reproduction, and the expression of ESBL, which can affect the structure of the peptidoglycan layer in the cell membrane. The cell growth was mathematically modeled using the generalized Verhulst equation. The parameter, characterizing the maximum possible number of cells grown under given conditions, decreased within the wild type (without plasmids) - p*bla*_TEM-135_ - p*bla*_TEM-1_ - p*bla*_TEM-20_ series, *i.e.*, the plasmid-bearing strains had reduced viability compared to that of the wild-type strain. The kinetics for the cell death of *N. gonorrhoeae* strains without the p*bla*_TEM-20_ plasmid in the presence of ceftriaxone can be described by a modified Chick-Watson law. For the *N. gonorrhoeae* strain that harbors the p*bla*_TEM-20_ plasmid, the kinetics for cell number changes with time reflected several processes: the hydrolysis of ceftriaxone by TEM-20 β-lactamase, growth and gradual death of cells. The demonstrated reduced viability of *N. gonorrhoeae* strains with the p*bla*_TEM-20_ plasmid probably explains the absence of clinical isolates of *N. gonorrhoeae* that produce ESBL.

## INTRODUCTION

The development of multidrug resistance in the *N. gonorrhoeae* pathogen is a major problem worldwide. According to the World Health Organization (WHO), gonorrhea may become incurable due to the ineffectiveness of old antimicrobial preparations (AMPs) and the lack of new AMPs for its treatment (Tacconelli et al., 2018; Golparian and Unemo, 2021).

Currently, and possibly in the future, antibiotics of the β-lactam group are the preferred drugs for treating gonococcal infection (Workowski and Bolan, 2015; WHO Guidelines, 2016; Tacconelli et al., 2018; Unemo et al., 2020). The group includes penicillin, cephalosporin, carbapenem, and monobactam subgroups. The mechanism of action of β-lactam antibiotics is to disrupt the synthesis of the peptidoglycan layer in bacterial cell walls, leading to bacterial death. In the mid-40s of the 20th century, penicillins started a revolution in the treatment of gonococcal infection and became the gold standard of treatment — a cure after a single injection. However, by the end of the 20th century, most clinical isolates of *N. gonorrhoeae* showed reduced susceptibility or resistance to penicillin, and for this reason, the use of penicillin was discontinued (Unemo and Shafer, 2014; Unemo and Jensen, 2017).

Now, for the treatment of gonococcal infections, the drugs of choice include another group of β-lactams, that is, third-generation cephalosporins, involving ceftriaxone, cefotaxime and cefixime, which are used as a monotherapy or in combination with the macrolide antibiotic azithromycin (Workowski and Bolan, 2015; WHO Guidelines, 2016; Tacconelli et al., 2018; Unemo et al., 2020). However, the levels of gonococcal resistance to cephalosporins are increasing every year, and these drug treatments were already reported to fail in several countries (Unemo et al., 2012; Allen et al., 2013; Golparian et al., 2014; Attram et al, 2019). The next group of β-lactams that could replace cephalosporins if cephalosporin resistance spreads is the carbapenem group (Unemo et al., 2020).

The resistance of *N. gonorrhoeae* to β-lactam antibiotics is associated with both chromosomal and plasmid determinants. Chromosomal mutations include substitutions in penicillin-binding proteins 1 and 2 (PBP1 and PBP2), mutations in the porin protein PorB that lead to a change in cell membrane permeability, and mutations that cause an increase in the expression level of MtrCDE efflux pump (Unemo and Shafer, 2014; Unemo and Jensen, 2017; Unemo et al., 2019). In this resistance phenomenon, the presence of the β-lactamase enzyme, which hydrolyzes the C-N bond in the β-lactam ring of the antibiotic and inactivates the action of AMP, is particularly important. The proportion of *N. gonorrhoeae* clinical isolates carrying p*bla*_TEM_ plasmids is not very high; for example, in the Russian population, the proportion remains at ~5% of the total number of isolates (Kubanov et al., 2019). However, the presence of β-lactamase in gonococci causes a significant increase in the level of resistance to penicillins (MIC_pen_ ≥ 16 mg/L) compared to chromosomal mutations (MIC_pen_ = 0.12-1.0 mg/L) (Shaskolskiy et al., 2019). To date, clinical isolates of *N. gonorrhoeae* have produced only enzyme variants that belong to broad-spectrum β-lactamases (penicillinases), which are unable to hydrolyze cephalosporins or carbapenems.

The following types of p*bla*_TEM_ plasmids are known for gonococci: Asian (7426 bp), African (5599 bp), Toronto/Rio (5154 bp), Nimes (6798 bp), New Zealand (9309 bp), Johannesburg (4865 bp) and Australian (3269 bp) (Müller et al., 2011). The plasmid *bla* gene has a length of 861 bp. Most penicillinase-producing isolates of *N. gonorrhoeae* contain the plasmid with the *bla*_TEM-1_ gene, but recently, the *bla*_TEM-135_ variant has been increasingly found in the worldwide population of *N. gonorrhoeae* (Ohnishi et al., 2010; Nakayama et al., 2012 Muhammad et al., 2014; Cole et al., 2015; Tanaka et al., 2021). The *bla*_TEM-1_ gene was found in both African and Asian plasmids, while *bla*_TEM-135_ was present mainly in Toronto/Rio and Asian type plasmids (Cole et al., 2015; Yan et al., 2019). *bla*_TEM-135_ differs from *bla*_TEM-1_ in one thymine-to-cytosine substitution at codon 182 (ATG→ACG). The resulting Met182Thr mutation is located far from the enzymatic active site (17 Å) in the hinge region between two β-lactamase domains and leads to stabilization of the β-lactamase structure (Orencia et al., 2001). As previously noted (Cole, 2015; Yan et al., 2019), the increased enzymatic stability may have contributed to the fixation of the TEM-135 β-lactamase variant in the *N. gonorrhoeae* population.

There is a large group of enzymes that are related to bacterial β-lactamases. TEM-1 and TEM-135 belong to class A serine β-lactamases according to the Ambler classification (Bradford, 2001; Bush and Jacoby, 2010; Tooke et al., 2019). This class also includes extended-spectrum β-lactamases (ESBLs), such as TEM-20, which are capable of hydrolyzing both penicillins and cephalosporins. Just one GGT→AGT nucleotide substitution in the *bla*_TEM-135_ gene, which results in the Gly238Ser mutation, enables gonococci to express TEM-20 type ESBLs (Arlet et al., 1999; Bradford, 2001). The TEM-20 β-lactamase variant already exists in nature. For example, in *Escherichia coli* (Adator et al., 2020), this variant resulted in increased MIC levels of cephalosporins (Jeong et al., 2004). The Gly238Ser substitution is associated with a change in the β-lactamase conformation, which increases the flexibility of the enzyme active site, and as a result, β-lactamase gains the ability to bind cephalosporins without losing its penicillin-hydrolyzing character (Orencia et al., 2001; Yan et al., 2019). Due to the increasing use of carbapenems in medicine, β-lactamases have emerged that can hydrolyze carbapenems; for example, Ambler class A *Klebsiella pneumoniae* carbapenemase (KPC) and Guiana extended-spectrum β-lactamase (GES) can be both chromosomal- and plasmid-localized (Tooke et al., 2019; Sawa et al., 2020). Thus, there is a risk that ESBLs may appear in gonococci, which could end the use of third-generation cephalosporins as treatments.

The goal of this work was to construct p*bla*_TEM_ plasmids that contain different variants of the *bla* gene, to prepare genetically engineered *N. gonorrhoeae* strains that contain p*bla*_TEM_ plasmids, to study the properties of *N. gonorrhoeae* strains that are transformed with the plasmids, and to assess the viability and resistance of these strains to β-lactam antibiotics (penicillins, cephalosporins and carbapenems). Particular attention was placed on constructing the p*bla*_TEM-20_ plasmid and studying the properties of the *N. gonorrhoeae* strain carrying this plasmid, which was not previously found in nature. This research will allow for the assessment of risks that are associated with the emergence and spread of third-generation cephalosporin-resistant strains and the possibility of further using β-lactam drugs for the treatment of gonococcal infection.

## MATERIALS AND METHODS

All experiments with *N. gonorrhoeae*, including transforming the cells, isolating plasmid DNA, electroporating the cells, cultivating the cells on solid media, washing the cells from Petri dishes and passaging them to other dishes, counting the grown colonies and assessing the number of viable bacteria (colony-forming units, CFU), were performed as described previously (Spence et al. al., 2008; Dillard, 2011).

### Isolation of the p*bla*_TEM-135_ plasmid from a clinical strain of *N. gonorrhoeae* with natural resistance to penicillins

We used an *N. gonorrhoeae* clinical isolate from the collection of the State Scientific Center of Dermatovenerology and Cosmetology of the Ministry of Health of the Russian Federation, obtained in 2017 from Chuvash Republic, and this isolate contained a Toronto/Rio type p*bla*_TEM-135_ plasmid, MIC_pen_ ≥ 32 mg/L (Shaskolskiy et al., 2019). The clinical isolate was seeded on GC chocolate agar (Thermo Fisher Scientific, USA) that was supplemented with IsoVitaleX Enrichment (Becton-Dickinson, USA) and contained 16 mg/L benzylpenicillin (Sigma–Aldrich, USA). The dish was incubated overnight at 37°C and 5% CO_2_. Grown colonies were harvested using a culture loop in 100 μl of PBS buffer. Isolation of plasmid DNA was performed with a Monarch Plasmid Miniprep Kit/T1010 (New England Biolabs, UK). The concentration and purity of DNA preparations were determined using a Nanodrop 2000 instrument (Thermo Fisher Scientific, USA). Plasmid DNA was analyzed by PCR as described previously (Palmer et al., 2000; Shaskolskiy et al., 2019).

### Construction of plasmid vectors with different variants of the *bla* gene on the basis of p*bla*_TEM-135_ from a clinical *N. gonorrhoeae* strain

Amplification of the *bla* gene fragment with simultaneous site-directed mutagenesis, which ensures that the mutations were introduced in the gene, was carried out. Using the primers 5′-TTACTTCTGACAACGATCGGAGGACCGAAGG-3′-FOR and 5′-AATGATACCGCGAGACCCACGCTCACTGGCT-3′-REV, a PCR fragment of the *bla* gene was obtained, and this fragment contained the GGT→AGT mutation that resulted in Gly238Ser and is present in TEM-20 ESBL (Figure 1). A PCR fragment corresponding to the part of the *bla* gene that encodes TEM-1 broad-spectrum β-lactamase was obtained in a similar way.

**FIGURE 1.**
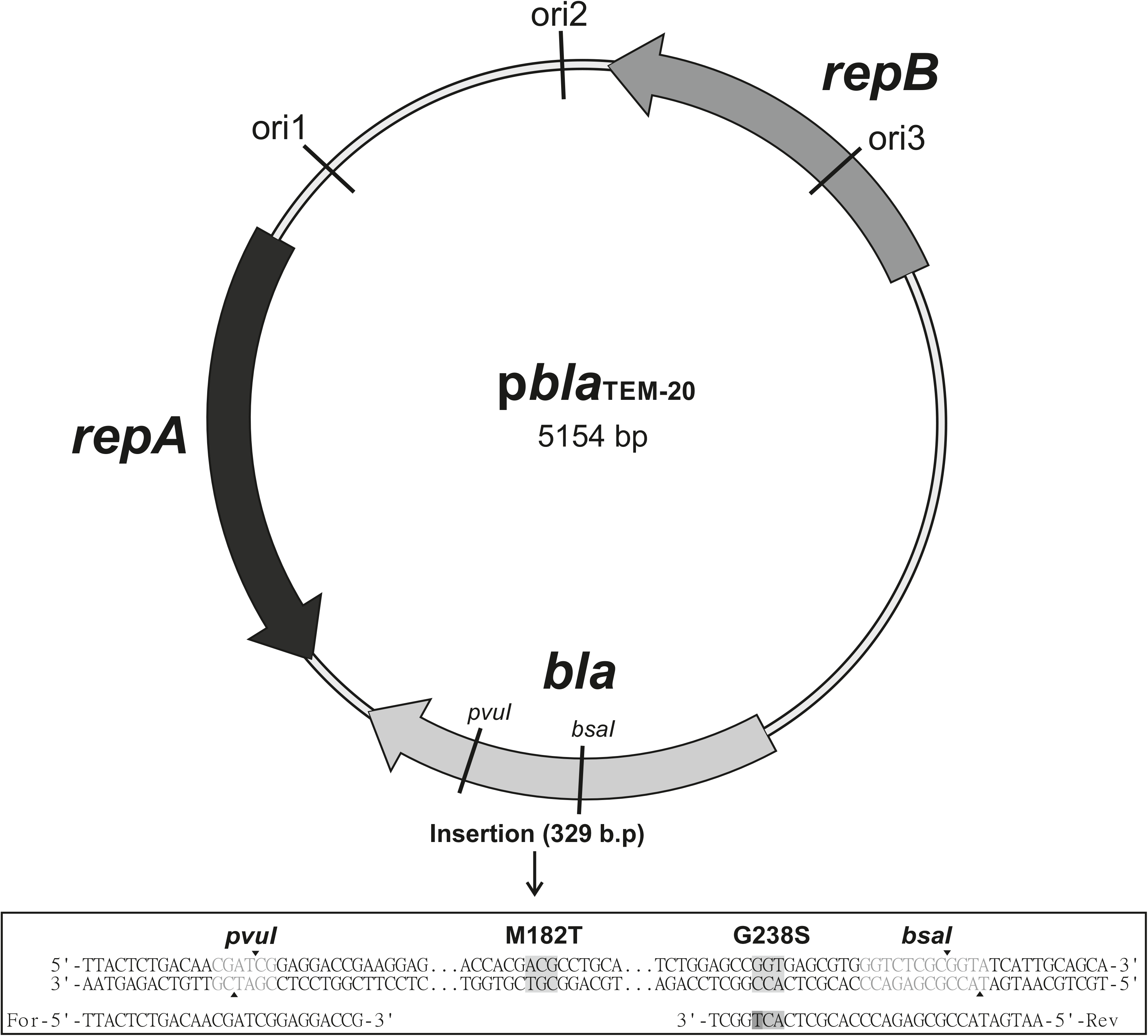
Map of the p*bla*_TEM-20_ plasmid vector, 5154 base pairs long, containing the *bla*_TEM-20_ gene associated with the resistance to β-lactam antibiotics, including penicillins and cephalosporins. The vector contained three replication origins (ori1, ori2, ori3) compatible with Gram-negative bacteria and the *repA* and *repB* genes encoding replication initiation proteins. Below is a nucleotide alignment of a 329 bp insert that contained a part of the *bla* gene with the GGT→AGT mutation, which resulted in a Gly→Ser substitution at codon 238, and the ATG→ACG mutation, which resulted in a Met→Thr substitution at codon 182.

The PCR fragments and the p*bla*_TEM-135_ plasmid were treated with the restriction endonucleases PvuI and BsaI. After that, the linearized p*bla*_TEM-135_ plasmid and the PCR fragments with sticky ends were treated with a ligation mixture to obtain the final p*bla*_TEM-1_ И p*bla*_TEM-20_ plasmids. The map of the resulting p*bla*_TEM-20_ plasmid vector is shown in Figure 1.

### Transformation of *E. coli* cells with plasmid vectors

The ready-to-use 10-beta competent *E. coli* strain DH10B (New England Biolabs, UK) was used for transformation with the plasmid vectors. The puc19 (K+) vector was used as a transformation control, and a sample without DNA was used as a negative control. The cells were grown and subsequently reseeded on selective media. Media containing 256-512 mg/L benzylpenicillin was used for *E. coli* with p*bla*_TEM-135_ И p*bla*_TEM-1_ plasmids; for cultures with p*bla*_TEM-20_ plasmid, the media contained ceftriaxone (Sigma–Aldrich, USA) at a concentration of 64 mg/L.

*E. coli* colonies grown on the selective media were collected with a culture loop in sodium phosphate buffer (PBS), and plasmid DNA was isolated using the Plasmid Miniprep/BC021S kit (Evrogen, Russia). The *bla* gene sequence and p*bla*_TEM_ plasmid type were confirmed by PCR and Sanger sequencing.

### Transformation of *N. gonorrhoeae* cells with plasmid vectors

We used *N. gonorrhoeae* cells, strain ATCC 49226 (F-18, CDC 10, 001, P935), which is a reference strain for antimicrobial susceptibility testing (https://www.culturecollections.org.uk/products/bacteria/detail.jsp?refId=NCTC+12700&collection=nctc). Transformation of *N. gonorrhoeae* with *pbla*_TEM-1_, p*bla*_TEM-135_ and p*bla*_TEM-20_ plasmids, which were extracted from *E. coli* cells, was performed by electroporation.

The optimal condition for transformation was resuspension of *N. gonorrhoeae* cells from the overnight culture inoculum, collected from the dish with a sterile culture loop, in 0.3 M sucrose. The cells were successively precipitated at 12000 RPM and resuspended in 0.3 M sucrose 2-3 times. For electroporation, 100-200 μl of purified cell suspension and 50-100 ng of the plasmid vector were put into 2 mm electroporation cells. A test tube without the addition of plasmid DNA was used as a negative control. Electroporation was performed by applying single pulsed discharges of 2.5 kV, 200 Ω, and 25 μF. The cooled cell suspensions were plated on chocolate agar dishes, washed and subcultured on dishes with a selective antibiotic; for strains with the p*bla*_TEM-1_ and p*bla*_TEM-135_ plasmids penicillin was used at a concentration of 2 mg/L, and for strains with the p*bla*_TEM-20_ plasmid ceftriaxone was used at a concentration of 0.5 mg/L.

The wild-type *N. gonorrhoeae* strain and strains with the *pbla*_TEM-1_ and p*bla*_TEM-135_ plasmids were stored at –80°C in cryo-medium containing trypticase-soy broth and glycerol at a ratio of 4:1.

The presence of β-lactamases in transformed cells was checked using nitrocefin discs (Remel, USA), which qualitatively detect the presence of enzymes, including penicillinases and cephalosporinases that are capable of destroying the β-lactam ring in the substrate (nitrocefin).

### Estimating the susceptibility of *N. gonorrhoeae* to β-lactam antibiotics

The following β-lactam antibiotics were used in this work: benzylpenicillin; ampicillin (aminopenicillin with the extended spectrum of action); cefuroxime (2-nd generation cephalosporin); ceftriaxone, cefixime and cefotaxime (3-rd generation cephalosporins); cefepime (4-th generation cephalosporin); and meropenem, imipenem, and doripenem (carbapenems), which were all from Sigma–Aldrich, USA.

The MICs of antibiotics for *N. gonorrhoeae* strains were measured by a serial dilution method. *N. gonorrhoeae* was cultivated on chocolate agar GC supplemented with IsoVitaleX Enrichment. An inoculum (0.5 McFarland) was prepared for the wild-type *N. gonorrhoeae* strain (ATCC 49226) and strains with the p*bla*_TEM_ plasmids. Microorganisms were seeded on Petri dishes containing selective medium with different concentrations of antimicrobial drugs. After incubation at 37°C with the addition of 5% CO_2_, the presence/absence of bacterial growth on Petri dishes was assessed. According to the measurement results, each strain was characterized in accordance with the established EUCAST criteria (https://www.eucast.org/clinical_breakpoints): S – susceptible, R – resistant.

### Assessment of *N. gonorrhoeae* cell viability: studies of cell growth and death

To estimate the effects of plasmids on the survival of gonococcus in the presence of antibiotics, the viability of *N. gonorrhoeae* cells that contain p*bla*_TEM_ plasmids was studied by obtaining cell growth curves in the absence of antibiotics and cell death curves in the presence of ceftriaxone. The change in the number of viable cells (colony-forming units, CFU) versus time was determined by counting the number of colonies grown on solid media.

All experiments during the studies of the growth and death of *N. gonorrhoeae* cells were performed immediately after transformation that allowed us to avoid the cell storage stage. The absence of storage was important because of the low viability of *N. gonorrhoeae* cells carrying the p*bla*_TEM-20_ plasmid.

To study cell growth, Petri dishes with chocolate agar (10 ml) without the addition of antimicrobials were used. An inoculum of cells (0.5 McFarland) was diluted to obtain ~50 CFU/ml, and 1 ml of cell suspension was plated on each dish, yielding 9 dishes that contained ~50 cells/dish for each *N. gonorrhoeae* strain. Every hour, the grown cells were washed off from every dish as follows: 1 ml of a sterile 0.3 M sucrose solution was poured onto the agar surface, the cells were resuspended on the dish with a sterile plastic loop without disturbing the agar surface layer, and the resulting cell suspension was transferred to a new dish. Thus, the cells grew on dish No. 1 for one hour, on dish No. 2 for two hours, and so on. The dishes with the washed-off cells were incubated for 24-48 hours at 37°C in the presence of 5% CO_2_, and the grown colonies were counted.

To study cell death in the presence of ceftriaxone, an inoculum of cells (0.5 McFarland) was diluted to obtain ~1000 CFU/ml, and 1 ml of cell suspension was plated on each dish with ceftriaxone, yielding 9 dishes that contained ~1000 cells/dish for each *N. gonorrhoeae* strain. Concentrations of ceftriaxone were 0.03, 0.125, and 2 mg/L. Every hour, the grown cells were washed off from every dish, and the resulting cell suspension was transferred to a new dish without the addition of ceftriaxone. The dishes were incubated as described above, and the grown colonies were counted.

### Mathematical models for cell growth and death

The generalized Verhulst equation was used to model cell growth curves for all strains under study (Peleg and Normand, 2007; Peleg and Corradini, 2011). To obtain cell death curves in the presence of ceftriaxone for the strains that did not contain the p*bla*_TEM-20_ plasmid, a modified Chick-Watson model (Jensen, 2010; Peleg, 2021) was used. The equation parameters were fit by enumerating the solutions of the Cauchy problem using MATLAB 2021b software for different *r*, *N_assymp_*, and α for the Verhulst equation and for different *k_obs_* values for the Chick-Watson equation. The optimization of the obtained numerical solutions was carried out by the least squares method when compared with the experimental results. The statistical significance of the difference between the mathematical models, which describe experimental results belonging to different data series, was evaluated by Fisher’s test (F test) in accordance with a procedure described previously (Motulsky and Ransnas, 1987).

## RESULTS

### Obtaining p*bla*_TEM-1_, p*bla*_TEM-135_, and p*bla*_TEM-20_ vector constructs

The p*bla*_TEM-135_ plasmid obtained from the clinical isolate of *N. gonorrhoeae* was used to construct artificial p*bla*_TEM-1_ and p*bla*_TEM-20_ vectors. As a result, three 5154 bp plasmids were obtained, which differed in the *bla* gene structure (Figure 1) as follows:

p*bla*_TEM-1_, no mutations in codons 182 and 238;
p*bla*_TEM-135_, ATG→ACG substitution at codon 182 leading to Met182Thr substitution, no mutation at codon 238;
p*bla*_TEM-20_, ATG→ACG substitution at codon 182 leading to Met182Thr substitution, and GGT→AGT substitution at codon 238 leading to the Gly238Ser substitution.

Plasmids could replicate in both *E. coli* and *N. gonorrhoeae* due to the presence of replication origins that were compatible with gram-negative bacteria and replication initiation proteins.

### Preparation of *E. coli* and *N. gonorrhoeae* strains containing p*bla*_TEM_ plasmids

*E. coli* and *N. gonorrhoeae* strains containing p*bla*_TEM-1_, p*bla*_TEM-135_, and p*bla*_TEM-20_ plasmids were obtained. *E. coli* strains with p*bla*_TEM_ plasmids were grown on selective media with antibiotics and stored on Petri dishes at 4°C.

The *N. gonorrhoeae* strains with p*bla*_TEM-1_ and p*bla*_TEM-135_ plasmids retained their viability after several (4-5) passages from dishes and after storage in cryo-medium. However, the *N. gonorrhoeae* strain with the p*bla*_TEM-20_ plasmid could not be stored in the cryo-medium, and the cells lost their viability after 12 hours of incubation on the plates; colonies did not grow after undergoing a subculture on the media both with and without antibiotics (penicillin, ceftriaxone). The loss of viability cannot be explained by the loss of plasmids, since the result of the test on nitrocephin disks was positive, indicating the presence of β-lactamases. PCR and Sanger sequencing also confirmed the presence of p*bla*_TEM_ plasmids with introduced mutations in the β-lactamase gene.

### Antimicrobial susceptibility testing of the *N. gonorrhoeae* strains containing different p*bla*_TEM_ variants

The susceptibility of *N. gonorrhoeae* strains that contain various p*bla*_TEM_ plasmids to several groups of β-lactam antibiotics was studied. The antibiotics included penicillins, which were previously used to treat gonococcal infection, cephalosporins, which are currently used for gonorrhea treatment, and carbapenems, which may replace cephalosporins if cephalosporin resistance spreads (Unemo et al., 2020). The MICs measured for these drugs are shown in Table 1.

**TABLE 1.** Susceptibility of *N. gonorrhoeae* strains containing different p*bla*_TEM_ plasmids to β-lactam antibiotics. S – susceptible, R – resistant, according to the EUCAST criteria (EUCAST, 2022)

| Antibiotic | EUCAST criteria | MIC, mg/L |  |  |  |
| --- | --- | --- | --- | --- | --- |
|  |  | WT* | *pbla*<sub>TEM-1</sub> | *pbla*<sub>TEM-135</sub> | *pbla*<sub>TEM-20</sub> |
| Benzylpenicillin (PEN) | S: MIC<sub>pen</sub> ≤ 0.06<br>R: MIC<sub>pen</sub> > 1 | 0.25 | 16 (R) | 32 (R) | 16 (R) |
| Ampicillin (AMP) | – | 0.125 | 8 | 8 | 16 |
| Cefuroxime (CXM) | – | 0.008 | 0.004 | 0.004 | 2 |
| Ceftriaxone (CRO) | S: MIC<sub>cro</sub> ≤ 0.125<br>R: MIC<sub>cro</sub> > 0.125 | 0.03 (S) | 0.015 (S) | 0.03 (S) | 4 (R) |
| Cefixime (CFM) | S: MIC<sub>cfm</sub> ≤ 0.125<br>R: MIC<sub>cfm</sub> > 0.125 | 0.015 (S) | 0.015 (S) | 0.015 (S) | 16 (R) |
| Cefotaxime (CTX) | S: MIC<sub>ctx</sub> ≤ 0.125<br>R: MIC<sub>ctx</sub> > 0.125 | 0.015 (S) | 0.008 (S) | 0.015 (S) | 8 (R) |
| Cefepime (FEP) | – | 0.03 | 0.015 | 0.015 | 16 |
| Meropenem (MEM) | – | 0.002 | 0.002 | 0.002 | 0.004 |
| Imipenem (IPM) | – | 0.004 | 0.008 | 0.004 | 0.008 |
| Doripenem (DOR) | – | 0.002 | 0.002 | 0.002 | 0.002 |

\*WT – wild-type, *N. gonorrhoeae*, ATCC 49226, without *pbla*<sub>TEM</sub> plasmid

After transformation of the wild-type *N. gonorrhoeae* strain with the p*bla*_TEM-1_ and p*bla*_TEM-135_ plasmid vectors, which contain genes encoding the broad-spectrum β-lactamase (penicillinase), the bacteria exhibited a sharp decrease in susceptibility to antibiotics from the penicillin subgroup (MIC_pen_ = 16-32 mg/L, MIC_amp_ = 8 mg/L). The MICs measured on the solid media confirmed that the TEM-1 and TEM-135 enzymes were unable to hydrolyze cephalosporins and carbapenems.

The MICs of the strain containing the p*bla*_TEM-20_ plasmid confirmed that the expressed TEM-20 β-lactamase is an ESBL that can hydrolyze both penicillins and cephalosporins of different generations. The cephalosporin MICs were above the EUCAST threshold for susceptible/resistant strains established to be 0.125 mg/L (Table 1). For example, the MICs of 3-rd and 4-th-generation cephalosporins were equal to or higher than 4 mg/L. Thus, the level of resistance of the strain with the p*bla*_TEM-20_ plasmid was higher than the threshold value established for 3-rd and 4-th-generation cephalosporins by more than a fivefold dilution of the antibiotic.

The measured MICs of carbapenems in all *N. gonorrhoeae* strains did not exceed 0.008 mg/L, *i.e.,* all strains were susceptible to carbapenems, proving that the TEM-1, TEM-135, and TEM-20 β-lactamase variants are unable to hydrolyze carbapenems (the EUCAST threshold for carbapenems is absent).

### Growth curves of the *N. gonorrhoeae* strains harboring different p*bla*_TEM_ plasmids, and the mathematical model for cell growth

Figure 2 shows the growth curves of the *N. gonorrhoeae* cells harboring the p*bla*_TEM-1_, p*bla*_TEM-135_, and p*bla*_TEM-20_ plasmids compared to that of the wild-type cells in the absence of antimicrobials. It should be noted that in our experiments we determined the number of viable cells calculated as the number of colonies grown on a dish (CFU). The wild-type strain and strains with p*bla*_TEM-1_ and p*bla*_TEM-135_ plasmids demonstrated bacterial growth during all 8 hours of cultivation. At the same time, the number of viable *N. gonorrhoeae* cells with the p*bla*_TEM-20_ plasmid began to decrease after 6 hours of cultivation.

**FIGURE 2.**
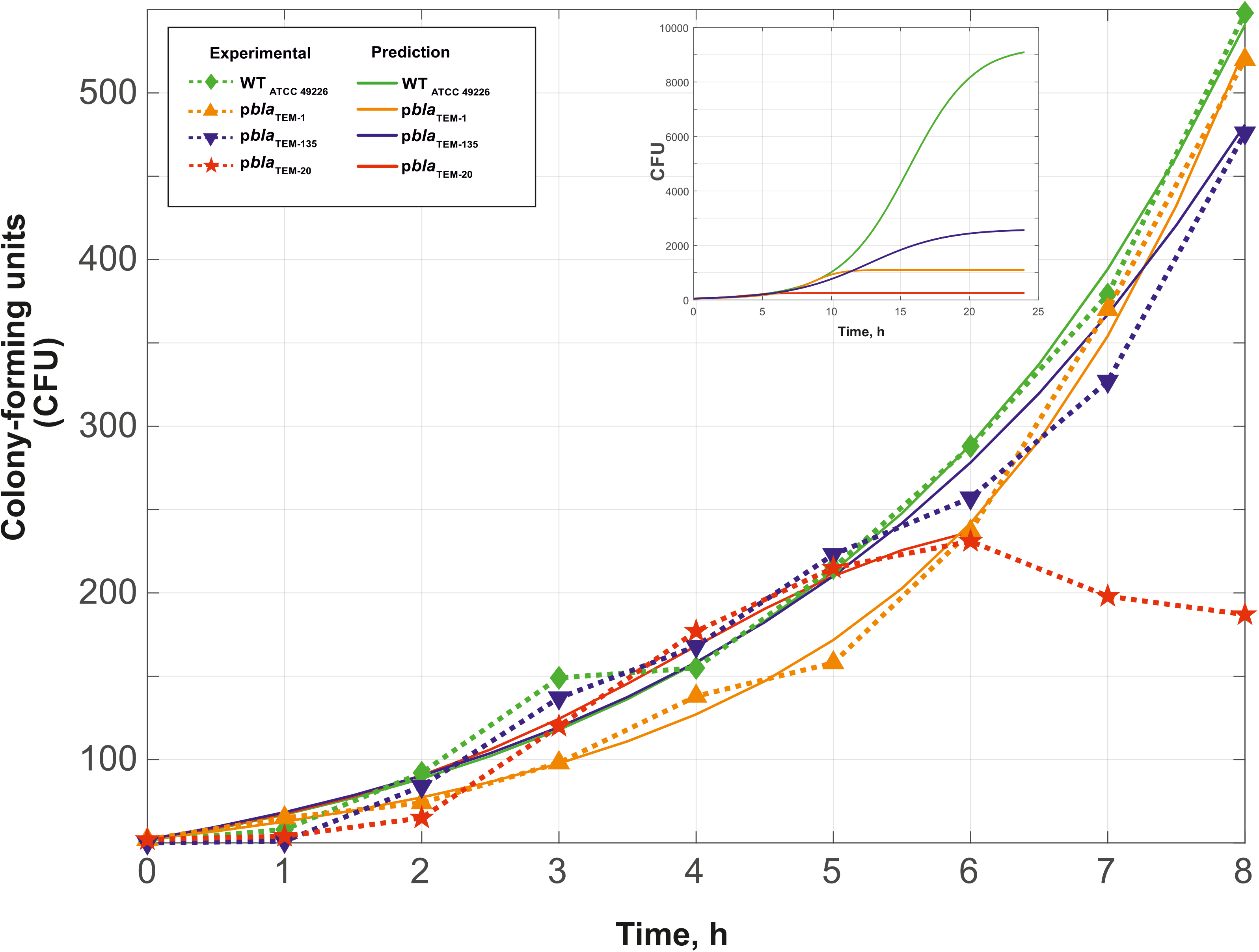
Growth curves of *N. gonorrhoeae* (change in CFU over time), strain ATCC 49226, without plasmids (WT) and with p*bla*_TEM_ plasmids, in the absence of antimicrobials (37°C, 5% CO_2_). Dashed lines and dots show the experimental results, and solid lines indicate the theoretical curves constructed using equations (2)–(5). The inset shows the theoretical curves up to 24 hours of incubation to reach *N*_*assymp*_.

To compare the *N. gonorrhoeae* strains that contain different plasmids, we performed mathematical modeling for the kinetic curves of cell growth. The generalized Verhulst equation (Peleg, 2007; Peleg, 2011) (eq. (1)) was chosen for modeling since it is the simplest and most universal equation for describing the dynamics of population growth up to the transition to the stationary phase.

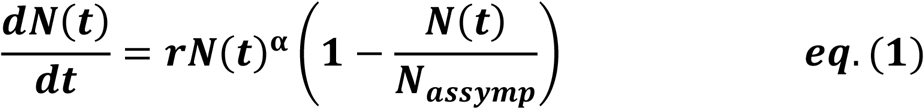

where *dN(t)/dt* is the growth rate at a given time;

*r* is the proportionality coefficient;

*N(t)* is the number of organisms at a given time;

*N*_*assymp*_ is the maximum possible number of grown cells of a given type under given experimental conditions;

α is the constant characterizing the growth peculiarity; α < 1 means that the organism does not achieve its potential for exponential growth; and α > 1 means that the organism exceeds this ability (Peleg, 2007).

Fitting of the parameters for equation (1) by enumerating the solutions of the Cauchy problems for different *r*, *N*_*assymp*_, and α resulted in equations (2)–(5), which describe the growth curves of cells with and without plasmids (Figure 2).

*N. gonorrhoeae* without plasmids (wild-type):

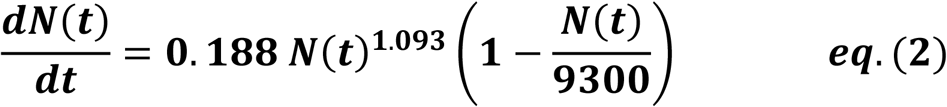

*N. gonorrhoeae* with p*bla*_TEM-1_:

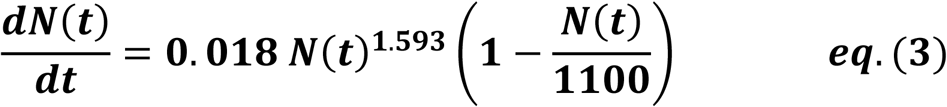

*N. gonorrhoeae* with p*bla*_TEM-135_:

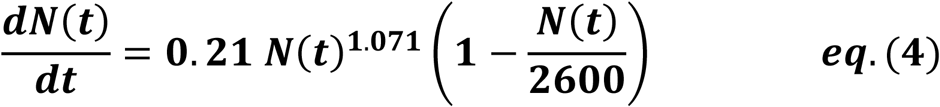

*N. gonorrhoeae* with p*bla*_TEM-20_ (up to 6 hours of growth):

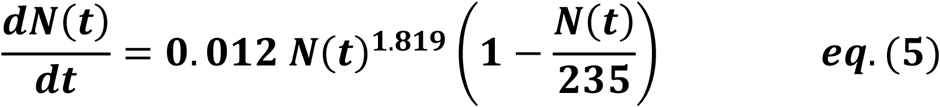

The R-squared coefficients were 0.9916 for the wild-type, 0.9958 for p*bla*_TEM-1_, 0.9801 for p*bla*_TEM-135_, and 0.9778 for p*bla*_TEM-20_.

The statistical significance of the difference between the obtained models for describing the data (*p* value) was calculated using the Fisher criterion. The wild-type model outperformed the model for TEM-1 with *p* = 0.02 and TEM-135 with *p* = 0.02. The model for describing TEM-1 was superior to that of the wild-type and TEM-135 models, with *p* = 0.01. The model for describing TEM-135 was superior to that of the wild-type model with *p* = 0.05 and the model for TEM-1 with *p* = 0.06. This indicates that the resulting models adequately describe cell growth for the corresponding strain.

The calculated value of *N*_*assymp*_ was different for the wild-type *N. gonorrhoeae* cells and for the cells with the p*bla*_TEM_ plasmids as follows: a maximum *N*_*assymp*_ of 9300 CFU was observed for the wild-type *N. gonorrhoeae*, lower values were observed for *N. gonorrhoeae* with the p*bla*_TEM-135_ and p*bla*_TEM-1_ plasmids, which were 2600 and 1100 CFU, respectively, and a very low CFU of 235 was observed for *N. gonorrhoeae* with the p*bla*_TEM-20_ plasmid. This means that the cells containing p*bla*_TEM_ plasmids had a reduced capacity of growth compared to that of the cells without plasmids, which can be explained by additional energy and metabolic costs for plasmid reproduction. At the same time, the strain with the p*bla*_TEM-20_ plasmid that expressed β-lactamase with the Gly238Ser substitution showed significantly reduced growth ability compared to that of the wild-type strain and strains harboring the other plasmids. The reduced viability of the strain with the p*bla*_TEM-20_ plasmid was also confirmed by the previously noted strain characteristics: the cells survived for no more than 12 hours when stored on solid media and did not survive when stored in a cryogenic medium.

### Change in the number of viable *N. gonorrhoeae* cells containing different p*bla*_TEM_ plasmids in the presence of ceftriaxone

Changes in the number of viable cells over time in the presence of various concentrations of ceftriaxone (0.03, 0.125 and 2 mg/L) are shown in Figure 3. Similar to the studies of bacterial growth, the CFU number on the solid media (chocolate agar) under various cultivation conditions was determined rather than the total number of cells. As shown in Figure 3A–C, the strains that did not contain ceftriaxone resistance determinants rapidly lost viability in the presence of this antimicrobial agent (the ceftriaxone concentrations in this experiment were equal to or greater than the MIC_cro_ for these strains, which were 0.015-0.03 mg/L (Table 1)). After 2-8 hours of incubation, the bacteria were completely extinct.

**FIGURE 3.**
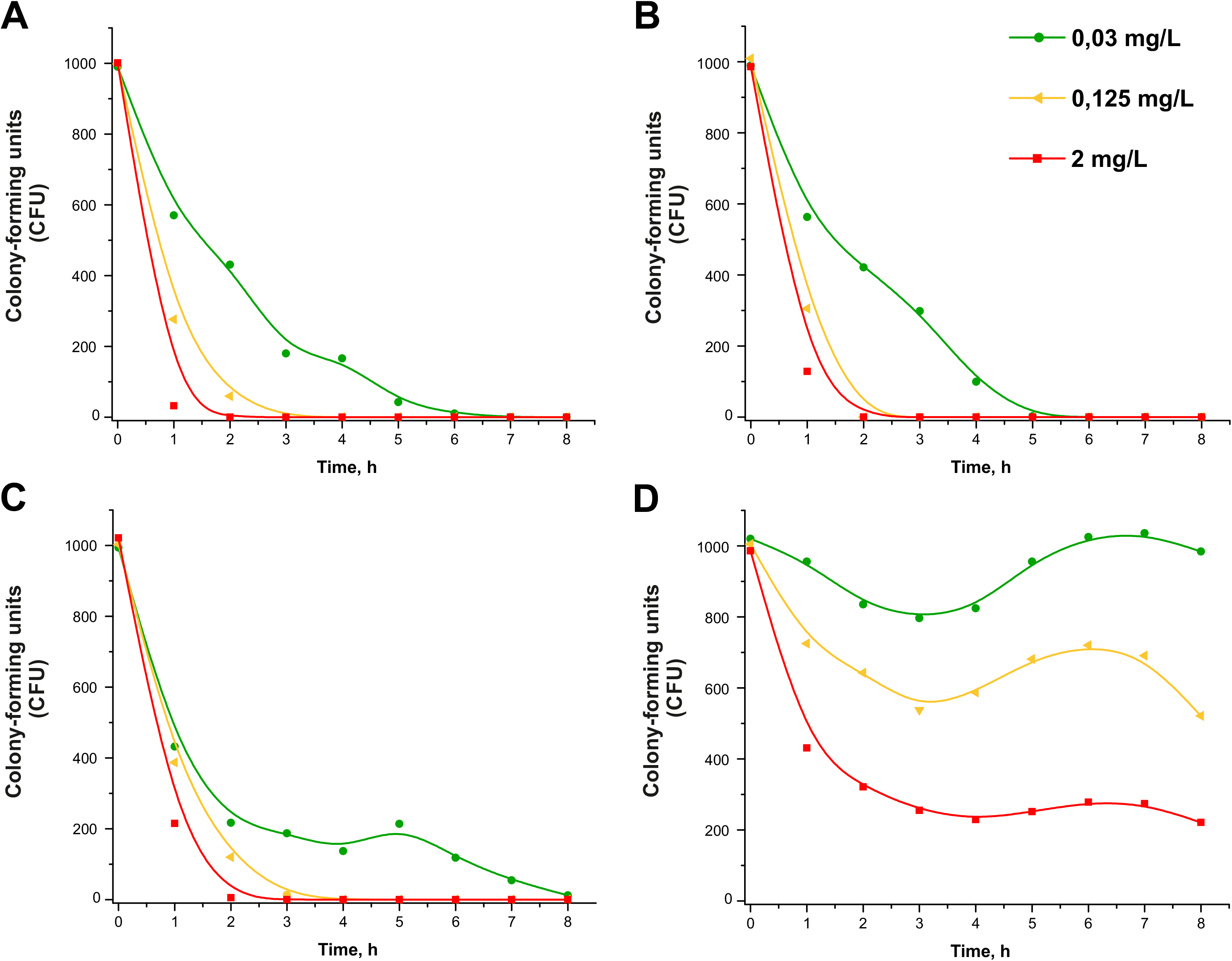
Change in the CFU number of *N. gonorrhoeae* in the presence of ceftriaxone 0.03; 0.125; 2 mg/L. *N. gonorrhoeae* strain ATCC 49226 without the plasmid (A), with p*bla*_TEM-1_ (B), p*bla*_TEM-135_ (C), and p*bla*_TEM-20_ (D) plasmids (37°C, 5% CO_2_).

Under our experimental conditions, the change in the concentration of ceftriaxone over time can be neglected, since this concentration is much higher than that of the cells in the sample; [CRO] = 0.03-2.0 mg/L or (0.045-3.030).10^−6^ M, *N*_0_ = 1000 cells (cells at zero time point). The decrease in CFU versus time for the curves in Figure 3A–C (strains without plasmids and with p*bla*_TEM-1_ and p*bla*_TEM-135_) followed a logarithmic law, *i.e.,* kinetic curves can be considered applicable for chemical reactions of the first (pseudo-first) order (Scheme 1).

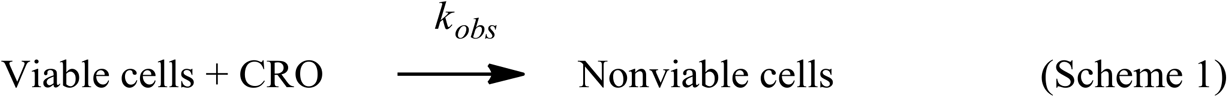

To describe the kinetic curves, a modified Chick-Watson model, which was developed for disinfection curves (cell death under the action of a disinfectant agent), can be applied. According to Chick’s law, the dependence of cell survival on time is described by the equations for the first-order chemical reaction (Jensen, 2010; Peleg, 2021) (equations (6) and (7)).

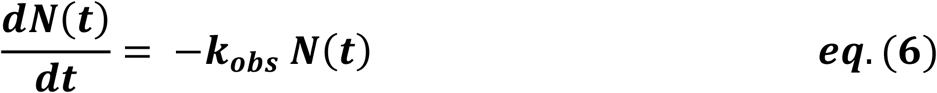

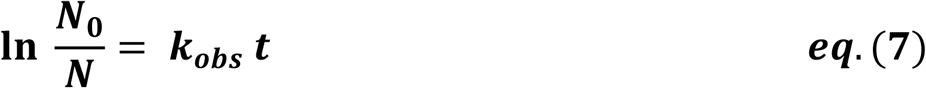

where *dN(t)/dt* is the growth rate at a given time;

*N(t)* is the number of cells at a given time;

*k*_*obs*_ is the observed first-order rate constant.

Fitting the cell death curves (Figure 3A–C) using equation (7) made it possible to obtain the values of the observed first-order rate constants *k*_*obs*_ for all strains at different concentrations of the disinfectant agent, which in our case was ceftriaxone (pseudo-first order rate constants). The dependences of *k*_*obs*_ on the concentration of ceftriaxone are shown in Figure 4.

**FIGURE 4.**
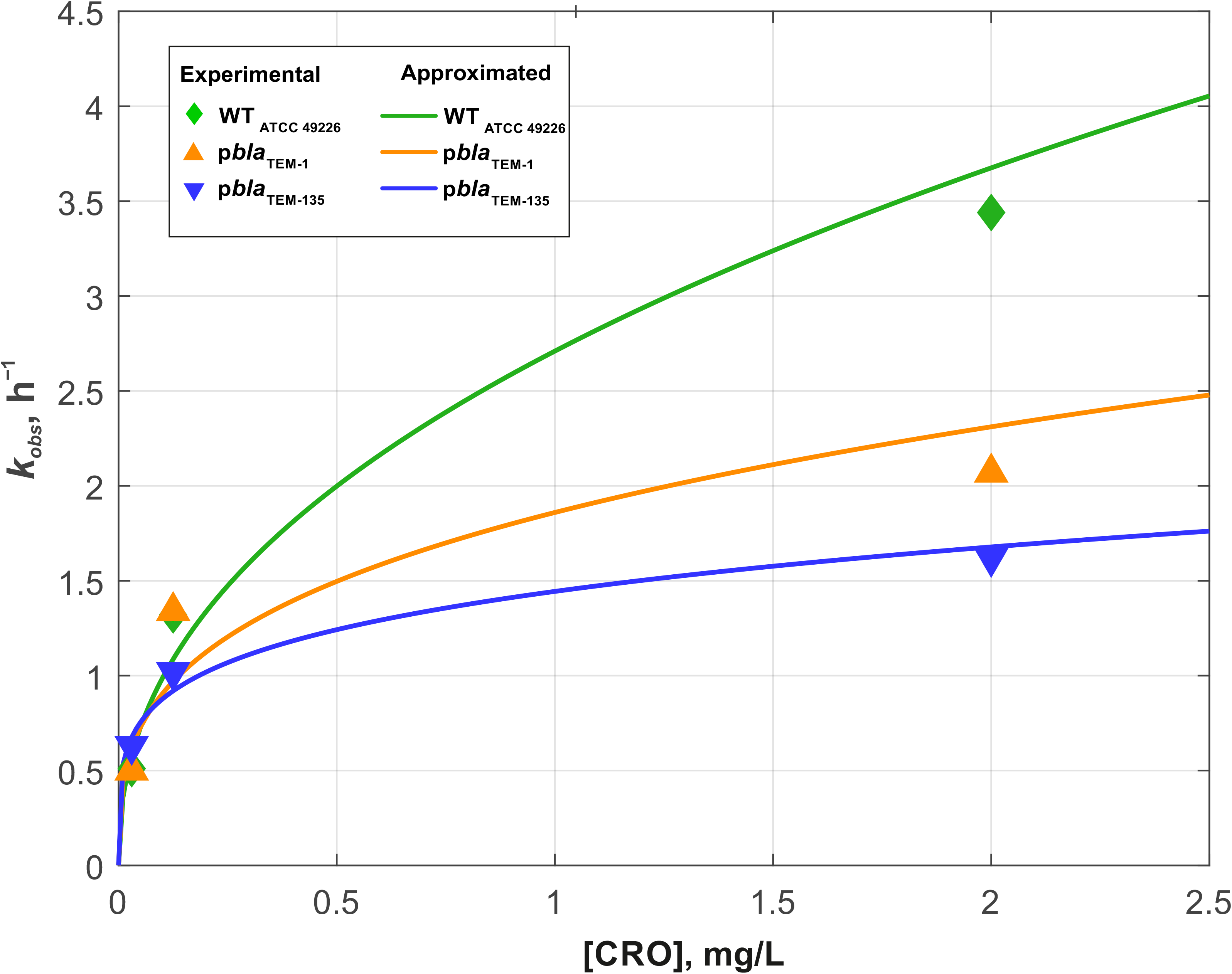
Dependences of the cell death rate constant *k*_*obs*_ on ceftriaxone concentration for *N. gonorrhoeae* strains without the plasmids and with p*bla*_TEM-1_ and p*bla*_TEM-135_ plasmids. The solid lines show the power-approximating curves constructed according to equations (10)–(12).

According to the modified Chick-Watson model, the dependence of the cell death rate on the concentration of ceftriaxone is determined by the following equation:

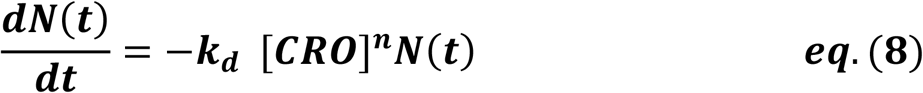

where [*CRO*] is the ceftriaxone concentration;

*n* is the fitting coefficient, which is also called the coefficient of dilution; and *k*_*d*_ is the true rate constant of cell death.

Thus, the observed death rate constant is related to the true rate constant and ceftriaxone concentration as follows:

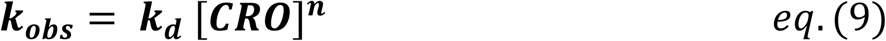

The dependences of *k*_*obs*_ on the concentration of ceftriaxone, shown in Figure 4, were approximated by the following power functions (R^2^ – approximation confidence value):

*N. gonorrhoeae* without plasmids (wild-type):

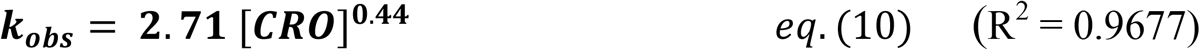

*N. gonorrhoeae* with p*bla*_TEM-1_:

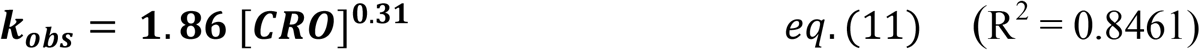

*N. gonorrhoeae* with p*bla*_TEM-135_:

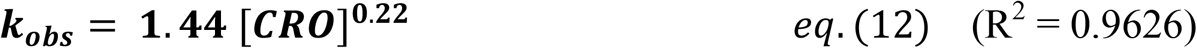

Thus, the equations describing the decrease in *N. gonorrhoeae* CFU versus time under the influence of ceftriaxone were as follows:

*N. gonorrhoeae* without plasmids (wild-type):

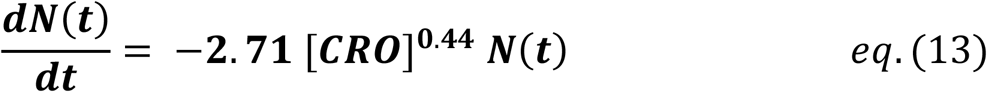

*N. gonorrhoeae* with p*bla*_TEM-1_:

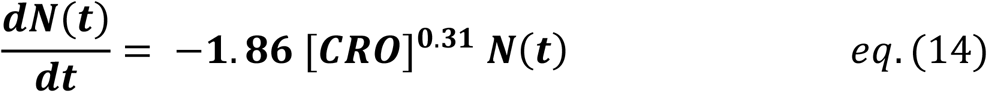

*N. gonorrhoeae* with p*bla*_TEM-135_:

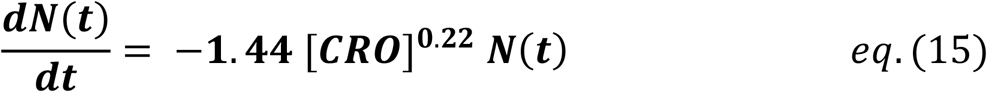

The results showed that the death of *N. gonorrhoeae* cells in the presence of ceftriaxone occurred differently in various strains. In the wild-type strain without p*bla*_TEM_, the cells died somewhat faster than in the strains carrying p*bla*_TEM-1_ and p*bla*_TEM-135_, although the β-lactamases expressed by these plasmids were not capable of destroying cephalosporins.

For the *N. gonorrhoeae* strain carrying the p*bla*_TEM-20_ plasmid that expresses ESBL, the CFU reduction versus time curves looked completely different (Figure 3D). The MIC of ceftriaxone for this strain was 4 mg/L (Table 1), and at a ceftriaxone concentration of 0.03-0.125 mg/L, some decrease in the number of live cells was observed in the first 3 hours. Then, a short-term increase in CFU was observed over a period of 3-6 hours, and this was changed to a reduction in CFU that was similar to the reduction observed in the *N. gonorrhoeae* strain with the p*bla*_TEM-20_ plasmid when it was cultivated in the absence of the antibiotic (Figure 2, curve for the strain with the p*bla*_TEM-20_ plasmid). At ceftriaxone concentration of 0.03 mg/L, *i.e.*, much less than the MIC_cro_ for the strain with p*bla*_TEM-20_, a significant amount of the antibiotic seemed to be hydrolyzed in the first 3 hours, and the number of cells reached its initial value during the next 3 hours. It should also be noted that the CFU versus time curves for this strain did not obey the logarithmic law, and in this case, the Chick-Watson model was not applicable to describe the kinetic curves.

Obviously, the kinetic curves for cells with p*bla*_TEM-20_ reflected several of the following processes: hydrolysis of ceftriaxone by TEM-20 β-lactamase, growth, and gradual cell death. Therefore, these curves could not be described by a simple Chick-Watson model for cell death in the presence of a disinfectant agent, which in our case was ceftriaxone.

## DISCUSSION

In this work, *N. gonorrhoeae* strains containing different types of β-lactamase plasmids were studied for their susceptibility to β-lactam antibiotics and for their viability. The artificial p*bla*_TEM-1_ and p*bla*_TEM-20_ plasmids were obtained by site-directed mutagenesis from the natural p*bla*_TEM-135_ plasmid of the Toronto/Rio type. Efficient plasmid amplification occurred in *E. coli* cells due to the presence in the vector’s structure of specific replication origins (ori1, ori2, ori3) and *repA* and *repB* genes encoding replication initiation proteins. Transformation of the *N. gonorrhoeae* strain ATCC 49226 with plasmid vectors was carried out using electroporation to obtain individual strains, for which the presence of p*bla*_TEM_ plasmids was confirmed. Among the strains, the *N. gonorrhoeae* strain was obtained with the p*bla*_TEM-20_ plasmid producing ESBL, which has not yet been found among clinical isolates.

Indeed, a single Gly238Ser amino acid substitution in the β-lactamase plasmid gene was sufficient to provide this enzyme with the ability to hydrolyze cephalosporins, while retaining the ability to destroy penicillins. However, the *N. gonorrhoeae* strain that contained the p*bla*_TEM-20_ plasmid showed substantially reduced viability; cell growth reached a plateau after 6 hours of cultivation both in the absence and in the presence of the antibiotic. In addition, the cells did not grow after incubation on plates for more than 12 hours and after freezing. Our results intersect with the work of Cole et al., 2018, in which genetically engineered strains of *N. gonorrhoeae* of the widespread NG-MAST type 1407, carrying African-type penicillinase-producing plasmids, were obtained. However, the strains described in the work of Cole et al. were unable to retain plasmids in the absence of antibiotic selection, and the cells lost plasmids even after one passage in the absence of penicillin. In our work, the reduced viability of the *N. gonorrhoeae* cells carrying the p*bla*_TEM-20_ plasmid was not associated with the loss of the plasmid during cultivation.

The reduced viability of the strain carrying the p*bla*_TEM-20_ plasmid that was demonstrated in this work can be explained by the following reasons: (a) the presence of the plasmid itself, which requires additional expenses from cells for its reproduction, and (b) the expression of ESBL. The action of cephalosporins, like all β-lactam antimicrobials, is aimed at inhibiting the synthesis of the bacterial cell wall by covalently inhibiting the action of transpeptidases (penicillin-binding proteins). The main component of the cell wall is peptidoglycan, which is a macromolecular structure that consists of peptide and sugar components. To protect themselves against β-lactam antimicrobials, bacteria express β-lactamases, which are located in the periplasmic space and hydrolyze the C-N bond in the β-lactam ring of the antibiotic, thereby inactivating the AMP. Because peptidoglycan in gram-negative bacteria possesses a C-terminal motif in the acyl-D-Ala-D-Ala peptide chain that is similar to the structural fragment of β-lactam preparations, β-lactamases can cause changes in the composition of peptidoglycan, thereby reducing the viability of bacteria. A change in the structure of the cell membrane leads to the suppression of cell division (bacteriostatic effect) or to the rupture of bacteria due to osmotic pressure (bactericidal activity) (Sawa et al., 2020). In the work of Fernández et al., 2012, it was shown that *E. coli*, which expressed certain β-lactamase variants, underwent a change in their peptidoglycan structures, in particular, a decrease in the level of cross-linked muropeptides that negatively affected the viability of such strains.

Evaluation of the resistance of the *N. gonorrhoeae* strains with p*bla*_TEM_ plasmids to β-lactam antibiotics also showed that none of the obtained β-lactamase variants, including ESBL TEM-20, were capable of hydrolyzing carbapenems. Thus, carbapenems remain among the drugs of the β-lactam series that can resist hydrolysis by *N. gonorrhoeae* bacterial lactamases, which confirms that they are potential drugs for the treatment of gonococcal infection.

The kinetic curves of cell growth were mathematically modeled using the generalized Verhulst equation (Peleg, 2007; Peleg, 2011). The value of *N*_*assymp*_, a parameter that characterizes the maximum possible number of cells grown under given conditions, was lower for the strains carrying the p*bla*_TEM_ plasmids compared with the cells without plasmids, *i.e.,* plasmid-bearing strains had reduced viability compared to that of the wild-type *N. gonorrhoeae*. *N*_*assymp*_ decreased within the series WT – p*bla*_TEM-135_ – p*bla*_TEM-1_ – p*bla*_TEM-20_. The reduced viability of the strains with p*bla*_TEM_ plasmids may explain why these strains have a lower frequency of occurrence in the *N. gonorrhoeae* population compared to that of the strains without plasmids. The higher viability of *N. gonorrhoeae* with p*bla*_TEM-135_ compared to that of *N. gonorrhoeae* with p*bla*_TEM-1_ may be associated with the presence of the Met182Thr mutation in β-lactamase, which leads to the stabilization of the structure of the enzyme. As noted previously (Orencia et al., 2001; Cole et al., 2015; Yan et al., 2019), increased enzyme stability may have contributed to the fixation and spread of the TEM-135 β-lactamase variant in the *N. gonorrhoeae* population.

The decrease in the number of viable cells with cultivation time in the presence of ceftriaxone has been studied. The kinetics of *N. gonorrhoeae* cell death without plasmids and with the p*bla*_TEM-1_ and p*bla*_TEM-135_ plasmids that lack ceftriaxone resistance determinants could be described by a modified Chick-Watson law for cell death modeling in the presence of a disinfectant (Jensen, 2010; Peleg, 2021). The death rate for the wild-type *N. gonorrhoeae* strain in the presence of ceftriaxone was higher than that of the strains with the p*bla*_TEM-1_ and p*bla*_TEM-135_ plasmids, although the strains carrying these plasmids were susceptible to ceftriaxone according to the EUCAST criteria.

Curves of the CFU versus time for the *N. gonorrhoeae* strain carrying the p*bla*_TEM-20_ plasmid that expresses ESBL cannot be described by a simple Chick-Watson model. The kinetic death curves in this case reflected several processes: hydrolysis of ceftriaxone by TEM-20 β-lactamase, growth, and gradual cell death.

The obtained data on the reduced viability of *N. gonorrhoeae* strains with the p*bla*_TEM-20_ plasmid may explain the absence of *N. gonorrhoeae* clinical isolates producing ESBL that hydrolyze cephalosporins of various generations.

## CONFLICT OF INTEREST

The authors declare that the research was conducted in the absence of any commercial or financial relationships that could be construed as a potential conflict of interest.

## ETHICS STATEMENT

Ethical approval/written informed consent was not required for the study of animals/human participants in accordance with the local legislation and institutional requirements.

## AUTHOR CONTRIBUTIONS

I.K. performed the experiments, analyzed results, and wrote the manuscript; D.G. designed and supervised the project, wrote the manuscript, A.V., O.A. and J.S. carried out antimicrobial susceptibility testing; A.K. and V.S. supervised work with cell cultures; D.D. wrote the manuscript;, B.S. directed the project, performed mathematical modeling and wrote the manuscript. All authors contributed to the article and approved the submitted version.

## FUNDING

This work was supported by the Russian Science Foundation, grant number 17-75-20039 (plasmid construction, susceptibility testing, mathematical modeling, cell growth characteristics) and by the Ministry of Science and Higher Education of the Russian Federation to the EIMB Center for Precision Genome Editing and Genetic Technologies for Biomedicine under the Federal Research Program for Genetic Technologies Development for 2019-27, agreement number 075-15-2019-1660 (gene sequencing and analysis of sequence data). Work with cell cultures was performed according to the Ministry of Health of the Russian Federation assignment number 056-03-2021-124.

## Notes

### Competing Interest Statement

The authors have declared no competing interest.

